# Stability and plasticity of contextual modulation in the mouse visual cortex

**DOI:** 10.1101/069880

**Authors:** Adam Ranson

## Abstract

Activity of neurons in primary sensory cortex is shaped by visual and behavioural context. However the long-term stability of the influence of contextual factors in the mature cortex remains poorly understood. To investigate this we used 2-photon calcium imaging to track the influence of surround suppression and locomotion on individual neurons over 14 days. We found that highly active excitatory neurons and PV+ interneurons exhibited relatively stable modulation by visual context. Similarly most neurons exhibited a stable yet distinct degree modulation by locomotion. In contrast less active excitatory neurons exhibited plasticity in visual context influence resulting in increased suppression. These findings suggest that the mature visual cortex possesses stable subnetworks of neurons, differentiated by cell-type and activity level, which have distinctive and stable interactions with sensory and behavioural context, as well as other less active and more labile neurons which are sensitive to visual experience.

## Introduction

Processing of information by cortical primary sensory circuits is strongly dependent upon both sensory and behavioural context. For instance in the primary visual cortex (V1) stimuli which extend beyond the classical receptive field (RF) of excitatory neurons typically suppress responses to RF stimulation (Adesnik et al., 2013; Blakemore and Tobin, 1972; Gilbert et al., 1996). Conversely behavioural factors such as locomotion increase activity of V1 neurons relative to states of quiet wakefulness (Ayaz et al., 2013; Keller et al., 2012; Niell and Stryker, 2010; Saleem et al., 2013).

A number of overlapping circuit elements have been suggested to mediate the effects of sensory and behavioural context on visual cortical processing. These include subclasses of local interneurons (Adesnik et al., 2013; Fu et al., 2014; Pi et al., 2013), innervation from the basal forebrain (Fu et al., 2014; Lee et al., 2014), noradrenergic innervation from the locus coeruleus (Polack et al., 2013) and input from the motor cortex (Keller et al., 2012). A fundamental question about these contextual effects, irrespective of their origin, concerns the stability vs flexibility of their influence in shaping early sensory processing in the mature cortex. Considerable effort has been invested in exploring the developmental plasticity of a range of rodent V1 receptive field properties during early post-natal development (Gordon and Stryker, 1996; Ko et al., 2014; Pecka et al., 2014; Wang et al., 2010) as well as the degree of plasticity of ‘classical’ receptive field properties such as orientation tuning and ocular dominance in the adult brain (Andermann et al., 2010; Kerlin et al., 2010; Kreile et al., 2011; Lütcke et al., 2013; Rose et al., 2016). In contrast many questions remain unaddressed regarding the long-term stability of contextual influences on mature V1 neurons (Gilbert and Li, 2013; Lütcke et al., 2013). Key issues include whether behavioural or visual context modulation is a fixed property of individual V1 neurons or only of the population as a whole; whether the degree of stability of modulation varies between cell types; and whether mature V1 neurons undergo experience dependent plasticity of contextual influences as has been previously demonstrated for orientation selectivity and ocular dominance.

We used longitudinal 2-photon imaging to track the stability of sensitivity to sensory and behavioural context of putative excitatory pyramidal neurons and parvalbumin positive (PV+) interneurons over an interval of 14 days, allowing repeated measurements of these features from single neurons. We found that mature V1 exhibits both long term stability of contextual modulation in a subset of highly responsive neurons, as well as a capacity for significant plasticity of more weakly responding cells, with PV+ inhibitory neurons exhibiting less plasticity in their responses than putative excitatory neurons.

## Results

In order to measure the stability of contextual factors in modulating neurons in primary visual cortex, 2-photon imaging together with the genetically encoded calcium indicator GCaMP6S was used to record longitudinally from putative excitatory and PV+ inhibitory neurons in both stationary alert, and running animals (Figure 1A and 1B). This allowed a repeated measurement to be made of the stability of modulation by contextual factors from single neurons (Figure 1C, 1D and 1E). Expression of the genetically encoded calcium indicator GCaMP6S was driven using an AAV injected into the primary V1 at a site targeted using intrinsic signal imaging (Figure 1B). In order to assess long-term stability of modulatory effects at the single cell level, neural activity was measured in adult animals (aged P80-P95) during a baseline session and then again 14 days (Δ14d) later. Average population retinotopic preference of recorded neurons was determined using 30 deg circular drifting gratings before the first session, with subsequent orientation and size tuning stimuli centred at this preferred retinotopic location.

**Figure 1.**
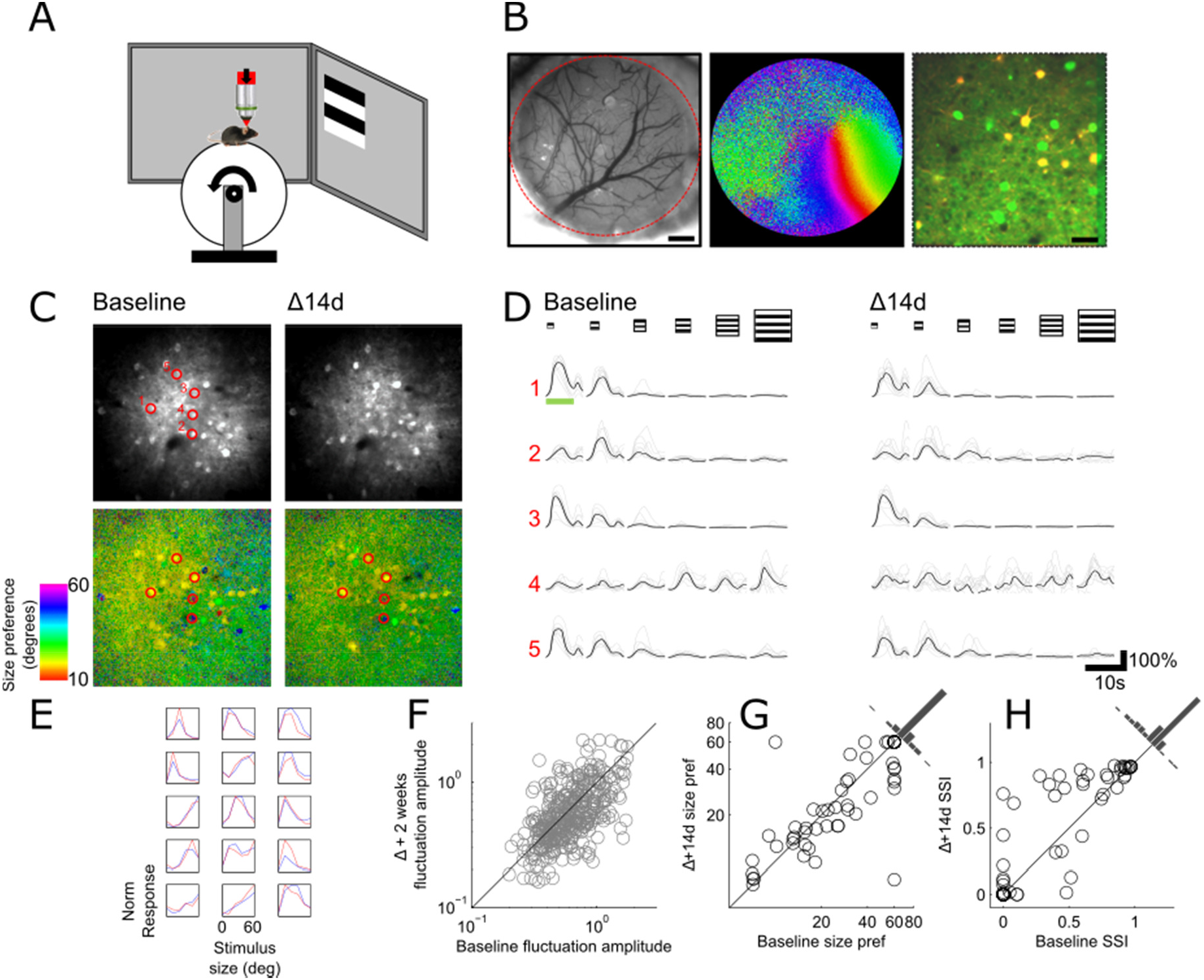
Experimental Setup and Longitudinal Response Stability. (A) Schematic of recording set up with mouse on fixed axis treadmill. (B) View through cranial window, showing cortical vasculature (scale bar = 500 µm), colour coded intrinsic signal imaging map of retinotopy, and co-localization image showing V1 neurons expressing GCaMP6S in all neurons, and tdTomato in PV+ cells (scale bar = 30 µm). (C) Representative example of field of view of V1 neurons recorded at baseline (top left) and 14 days later (top right). Pixel-wise stimulus preference map from same field of view showing size preference on a pixel-by-pixel basis at baseline and 14 days later (bottom left and right, scale bar = 30 µm). (D) Traces of 5 example neurons marked in (C) showing baseline and 14d later responses to size tuning stimulus, with light traces showing individual trials, and black traces showing average response. Green bar shows stimulation period. (E) Example size tuning curves of 18 highly responsive neurons showing stability of tuning between baseline (blue) and 14d later. (F) Between session correlation in magnitude of calcium signal fluctuations. (G-H) Between session correlation of size preference and surround suppression index. See also Supplementary Video 1.

### Highly responsive V1 neurons exhibit long-term stabiiity of size preference over time

For each field of view we first calculated pixel-wise maps of size preference in which a high degree of diversity of size preference could be observed (Figure 1C, bottom panels). Single neurons could easily be visually discriminated as clusters of pixels with similar preference, and many of these clusters could be seen to persist over the two sessions. In order to measure size preference of individual cells over the two sessions, regions of interest which correspond to individual neurons were detected using a semi-automated algorithm (see methods), and these were used to extract single cell responses over the two sessions. Size preference and surround suppression index (SSI, see methods) was measured for putative excitatory neurons which had an orientation preference close to the size tuning stimulus (+/-30°, horizontal grating) and exhibited robust visually evoked responses (ΔF/F > 1) which persisted over the 2 recording sessions. This highly responsive subset of cells was selected to minimise the influence of measurement noise in assessing the stability of surround suppression. Intersession correlation of activity fluctuations in the excitatory population as a whole (quantified as the standard deviation of each neurons ΔF/F trace) showed that highly responsive cells tended to be persistently highly responsive (r = 0.57, n = 383, p < 10^−33^, Figure 1F, Supplementary Figure 3C and 4). However despite this, only 47% of the top 20% most active neurons in the first session were also in this category in the second session, indicating significant motility in average activity levels. Persistently highly responsive cells spanned the full range of size preferences measured (Figure 1D, 1E and 1G), and exhibited a range of degrees of suppression (Figure 1D, 1E and 1H). There was however a bimodality in the distribution of both size preference and surround suppression index (SSI) whereby extremes were over represented (see Supplementary Figure 1). Comparison of size tuning curves between sessions revealed a high degree of inter-session correlation of both size preferences (r = 0.80, n = 60, p < 10^−13^, Figure 1G) and SSI (r = 0.86, n = 60, p < 10^−18^, Figure 1H) with a mean absolute difference of 6.9 ± 1.57° in size preferences. While size preference of individual cells did not differ significantly between sessions, there was a net increase in the degree of suppression of 0.08±0.03 (n = 60, p < 0.05). This analysis shows that a subpopulation of persistently highly responsive cells maintain robust size preferences over time, consistent with a scheme of lifetime sparseness (Barth and Poulet, 2012).

### Less active V1 neurons exhibit robust experience dependent plasticity of surround suppression

The population of excitatory neurons was next divided into 3 groups depending on response magnitude to preferred stimuli (ΔF/F low 0.3-0.7, n = 48, medium 0.7-1, n = 19, and high >1, n = 60) with a requirement imposed that neurons had to fall into the same category over the two sessions to be included in further analysis. This allowed an examination of whether stability of surround suppression varies depending upon overall responsiveness. While high responding cells were found to exhibit long-term stability in their degree of modulation by visual context (as described above), more weakly responding cells exhibited a higher degree of intersession variability in both size preference and SSI (Figure 2). However rather than simply reflecting greater measurement noise (e.g. due to lower response magnitudes, see Supplementary Figure 2 for within vs. between session analysis), or differences in behavioural state (see Supplementary Figure 2E, Supplementary Figure 6), this inter-session variability was instead due to near uniform systematic decreases in preferred size, and increase in SSI of the majority of the neurons recorded. In the population of excitatory neurons as a whole the majority of cells either decreased or maintained preferred size (81%), and increased or maintained SSI (80%) as can be observed in figure 2B and 2C in which most data points can be seen to be below and above the unity line respectively. The degree of net reduction in size preference was next quantified in the 3 response level groups (Figure 2D) and significant plasticity was observed in both low responding neurons (size shift = −18.08° ± 2.64°, n = 48, p < 10^−7^, paired t-test) and medium responding neurons (size shift = −7.52° ± 3.23°, n = 19, p < 0.05, paired t-test) but not high responding neurons (size shift = −2.31° ± 1.69°, n = 60, p = 0.18, paired t-test), as well as a small but statistically significant reduction is response amplitude in the low responding group (reduction in ΔF/F = 0.087, p<0.05, Supplementary Figure 3A and 4). A similar pattern emerged for SSI (Figure 2E) which was strongly increased in low responding neurons (SSI shift = 0.27 ± 0.04, n = 48, p < 10^−6^, paired t-test) and medium responding neurons (SSI shift = 0.29 ± 0.06, n = 19, p < 10^−3^, paired t-test), and less strongly but significantly increased in strongly responding neurons (SSI shift = 0.08 ± 0.03, n = 60, p < 0.05). These results indicate that while strongly responding neurons maintain stable size preferences over time and are relatively immune to the effects of visual experience, less active neurons can exhibit a high degree of plasticity, which can be recruited by visual experience, and results in a driving up of suppression and a driving down of preferred stimulus size.

**Figure 2.**
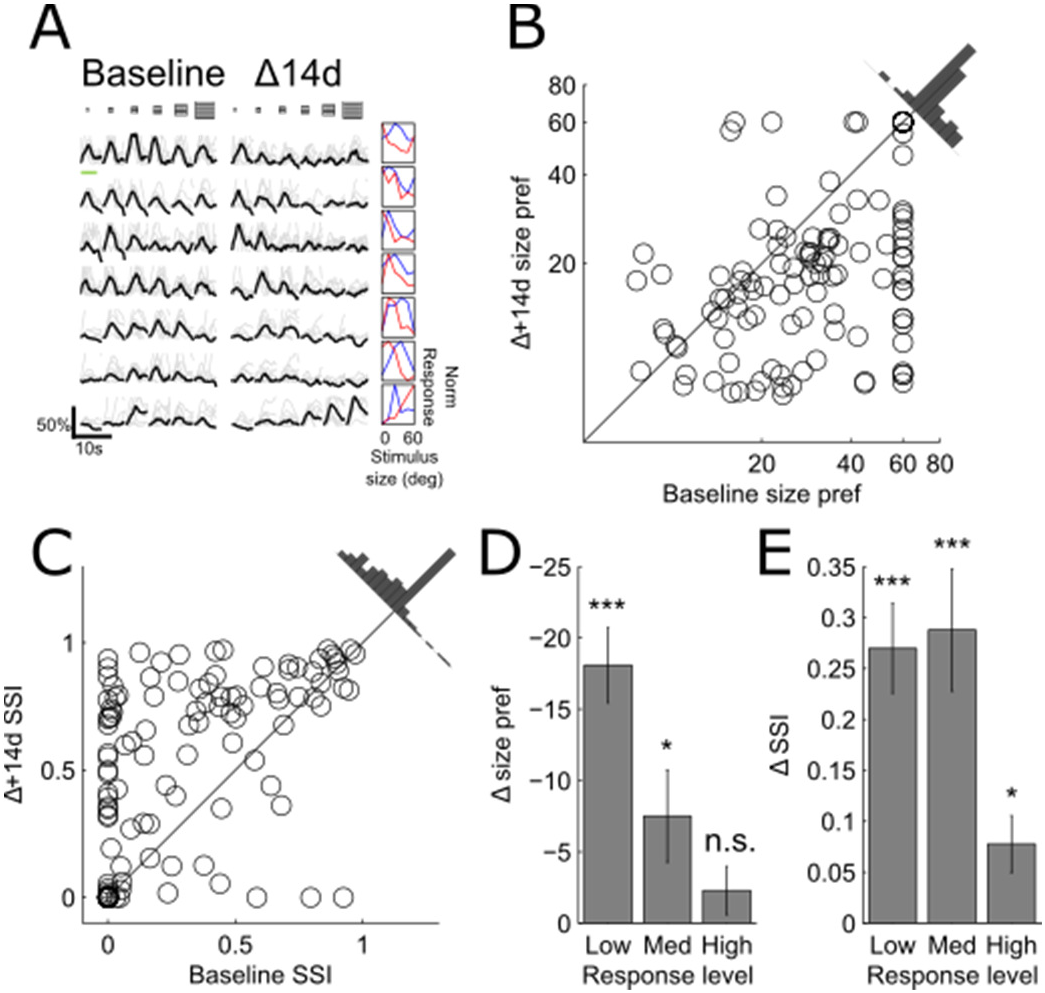
Plasticity Of Size Preference and Suppression in Weakly Responding Neurons. (A) Traces of 7 example neurons marked in showing baseline and 14d later responses to size tuning stimulus, with light traces showing individual trials, and black traces showing average response. Green bar shows stimulation period. Normalized raw tuning are shown on the right. (B-C) Intersession correlation of size preference (B) and SSI (C) showing shift in response preferences towards smaller sizes and greater suppression. (D-E) Average net shift in size preference (D) and SSI (E) in low medium and high responding neurons shows greater plasticity in weakly responding neurons. All data are presented as mean ± SEM ***p < 0.001; **p < 0.01; *p < 0.05; n.s., not significant.

### V1 neurons exhibit longitudinally stable and characteristic modulation of activity by locomotion

We next investigated the stability of the effects of locomotion on visual cortex activity. As has been previously reported, activity of excitatory neurons in V1 was strongly modulated by locomotion (Figure 3A and 3B). The correlation between putative excitatory neuron activity and locomotion was calculated in the population as a whole during ongoing visual stimulation with the size tuning stimulus. Differences between experiments in proportion of time moving vs. stationary introduce artefactual variability in locomotion-neural activity correlation. In order to mitigate this problem, data was subsampled to include equal periods of time of locomotion and stationary behaviour (see methods for detailed description and Supplementary Figure 5 for example cells and summary histograms). The activity of the vast majority of putative excitatory neurons was correlated with locomotion (Figure 3C), with significant correlation coefficients (p < 0.01, shuffle test) ranging from −0.30 to 0.81, and with a stable fraction of significantly correlated neurons observed over the 14d period (baseline = 88.8%, Δ14d = 89.2%). We next assessed whether the degree of locomotion modulation of individual putative excitatory neurons was stable over the two sessions. Correlating individual cell's baseline locomotion correlation with its Δ14d locomotion correlation, revealed a strong inter-session correspondence across the range of locomotion correlation values (r = 0.64, n = 383, p < 0.01; Figure 3C). This suggests that despite the proposed relatively indirect mechanisms of modulations of neural activity by locomotion (Fu et al., 2014; Lee et al., 2014), the modulation is in large part a stable feature of individual cells over time. In addition further examination of the low responding group, which exhibited increases in surround suppression between sessions, revealed that these neurons also exhibited a small but statistically significant reduction in correlation with locomotion (shift = −0.13, p < 0.01, t-test, Figure 2E) which was not present in more strongly responding neurons (shift = −0.02, p = 0.68, t-test). We next investigated whether the degree of modulation of neural activity by locomotion was a stable feature of individual neurons. The degree of locomotion induced gain of visual responses to preferred size stimuli was quantified with a locomotion modulation index (LMI) defined as (R_M_-R_S_)/R_S_ where R_M_ and R_S_ are the visual response while moving and stationary respectfully. LMI was compared between baseline and Δ14d sessions for the subset of neurons which had stationary and locomotion trials for their preferred stimulus size, and were either in the low responding group ΔF/F 0.3-0.7) or high responding ΔF/F >1). This analysis revealed that while there was a strong positive correlation between sessions in LMI in the population as a whole (r = 0.48, n = 49, p < 0.01, Figure 3D), there was in addition a systematic inter-session increase in LMI which was observed in the low responding group (shift in LMI = 1.04, p < 0.001, paired t-test, Figure 3F) but not in the high responding group (shift in LMI = 0.17, p = 0.15, paired t-test, Figure 3F), and in addition that this plasticity was limited to responses to the preferred stimulus (Figure 3F). These findings show that the majority of excitatory V1 cells are significantly modulated by locomotion, and that while this modulation varies significantly between neurons, is in large part stable over the 14d interval within single neurons. In addition this data indicates that in adult animals the modulation of visual response by locomotion is plastic.

**Figure 3.**
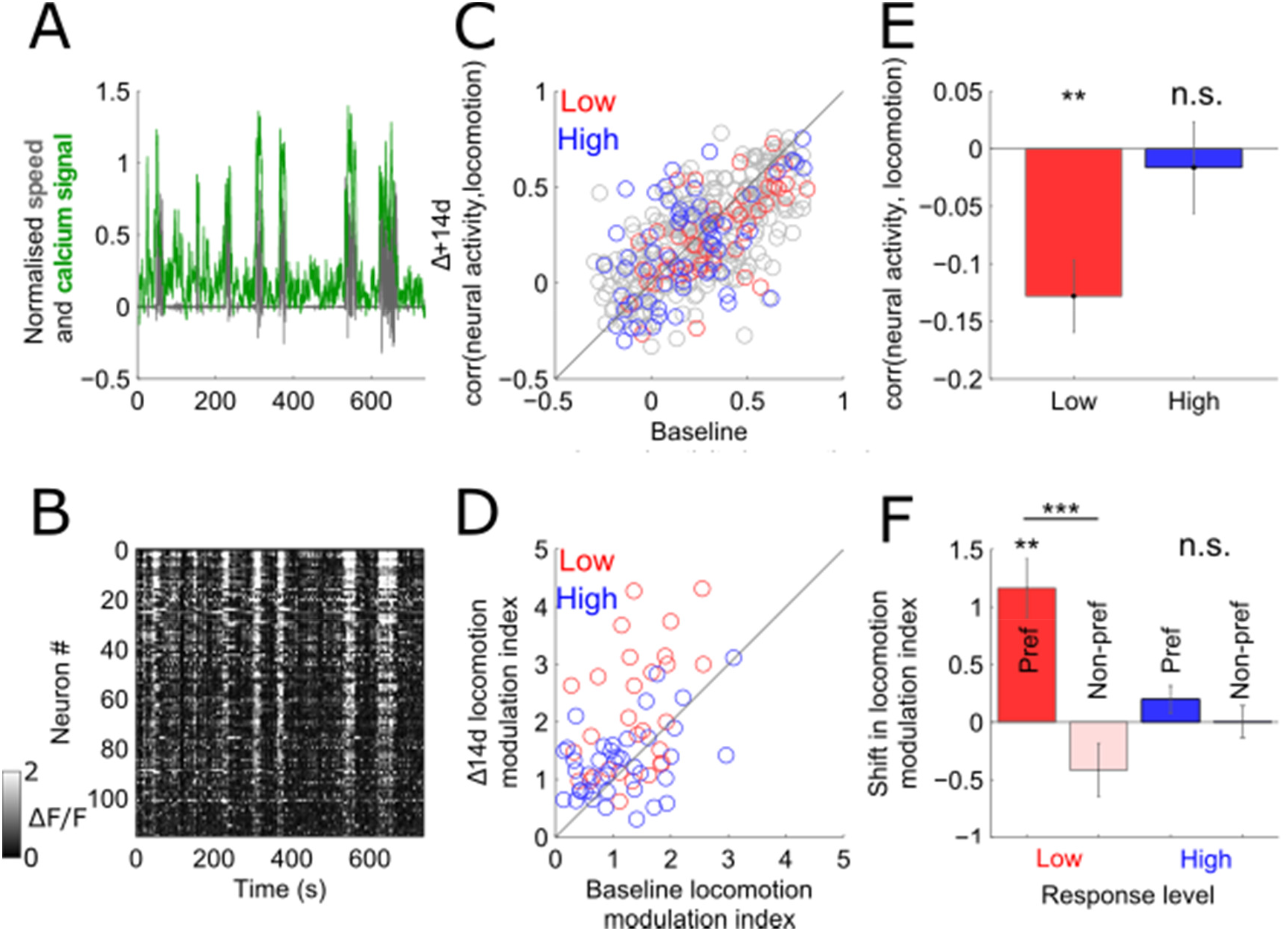
Stability of modulation of neural activity by locomotion. (A) Overlaid normalised traces of locomotion speed (red) and average neural activity of a population of 109 neurons, showing high degree of correlation between neural activity and locomotion. (B) Activity map of neurons averaged in (A) over the same time period. (C-D) Correlation of neural activity with locomotion (C) and modulation of visually evoked response to preferred stimulus (D) between baseline session verses session 14d later. (E-F) Between session shift in corr(neural activity, locomotion) (E) and LMI (F) in low and high responding neurons during preferred and non-preferred stimulus presentation. All data are presented as mean ± SEM ***p < 0.001; **p < 0.01; *p < 0.05; n.s., not significant.

### PV interneurons exhibit a greater degree of long-term stability of modulation by contextual factors than putative excitatory neurons

We next measured the stability of modulation of PV+ neurons (Figure 4A) by these contextual factors. Previous work has provided evidence that PV+ interneurons receive input from a relatively unbiased sample of local excitatory neurons (Cossell et al., 2015), a model which would predict that PV neurons would show relatively homogenous modulation by contextual factors, with parameters reflecting the average of the excitatory population as a whole (as has been shown with orientation selectivity). Contrary to this prediction, at baseline PV neurons exhibited heterogeneous size preferences and SSIs, and also exhibited a similar bimodal distribution of both size preference and SSI as putative excitatory neurons (Supplementary Figure 1C and 1D). In common with the putative excitatory population, more active neurons during the baseline session tended to be more active during the Δ14d session (Figure 4B). The intersession correlation of size preference and SSI were next calculated in the PV+ neuron population as a whole (using the inclusion criteria of maintaining a response amplitude across the two sessions of ΔF/F>0.3) and in the highly responding subset of cells ΔF/F>1). Size preferences and SSI were both highly correlated between sessions, both in the population as a whole (Size: r = 0.70, n = 123, p < 10^−18^; SSI: r = 0.71, n = 123, p < 10^−20^; Figure 4C and 4D) and in the highly responding subset of cells (Size: r = 0.88, n = 36, p < 10^−09^; SSI: r = 0.94, n = 30, p < 10^−13^; Figure 4C and 4D). We next tested for systematic between session plasticity of size tuning and SSI as had been observed in putative excitatory neurons, comparing plasticity in weakly and strongly responding cells using the same criteria employed in the excitatory population. In common with the putative excitatory population, strongly responding PV+ neurons exhibited no significant shift between sessions in size preference (mean shift = −2.62 ± 1.68, n = 36, p = 0.13; Figure 4E), and in addition exhibited no significant shift in SSI (mean shift = 0.05 ± 0.02, n = 36, p = 0.052; Figure 4F). In contrast to the excitatory population, weakly responding PV+ neurons exhibited no significant net shift in either preferred stimulus size (mean shift = −6.74 ± 3.32, n = 19, p = 0.06; Figure 4E) or SSI (mean shift = 0.12 ± 0.08, n = 19, p = 0.15; Figure 4F). A between cell type comparison of mean shift in size preference and SSI confirmed a significant difference in PV+ vs. PV-cells in between session plasticity (size shift: p < 0.01, t-test; SSI shift: p < 0.05, t-test). Despite the lack of statistical significance of shifts of size and SSI in PV+ cells, the degree of variability in the weakly responding population, particularly in the shift in SSI, suggests that there may be a subset of PV+ which are more plastic and exhibit a similar behaviour to weakly responding excitatory neurons, but which are not distinguishable by their response amplitude. The stability of modulation of PV+ neuron activity by locomotion was next measured. As was observed in excitatory neurons there was both a large fraction of PV+ cells with significant correlations with locomotion (91% in session 1, 95% in session 2, Figure 4G), and a high degree of intersession correlation in degree of modulation (r = 0.71, n = 196, p < 10^−30^). A between cell-type comparison of baseline neural activity correlation with locomotion revealed no significant difference between the 2 cell types (PV+ = 0.29 ± 0.02; putative excitatory = 0.269 ± 0.01; p = 0.32, t-test). We next compared the degree of stability of the modulation of visually evoked responses by locomotion in the low responding and high responding PV+ population. In common with the high responding putative excitatory neurons, high responding PV+ neurons exhibited no significant mean shift in LMI between sessions (shift = 0.18, n = 36, p = 0.11, paired t-test). Low responding neurons also exhibited no systematic shift which reached statistical significance (shift = 1.55, n= 19, p = 0.15) although in common with their shift in SSI there was a high degree of diversity in shift size within this group. These results indicate that the responses of PV+ interneurons are relatively stable in general compared to the excitatory population, especially in a highly responsive subpopulation, despite significant shifts in the local excitatory population. In particular the PV+ population did not undergo the robust between session increase in surround suppression, and reduction in preferred stimulus size observed in excitatory neurons, although it did exhibit a weak trend in this direction.

**Figure 4.**
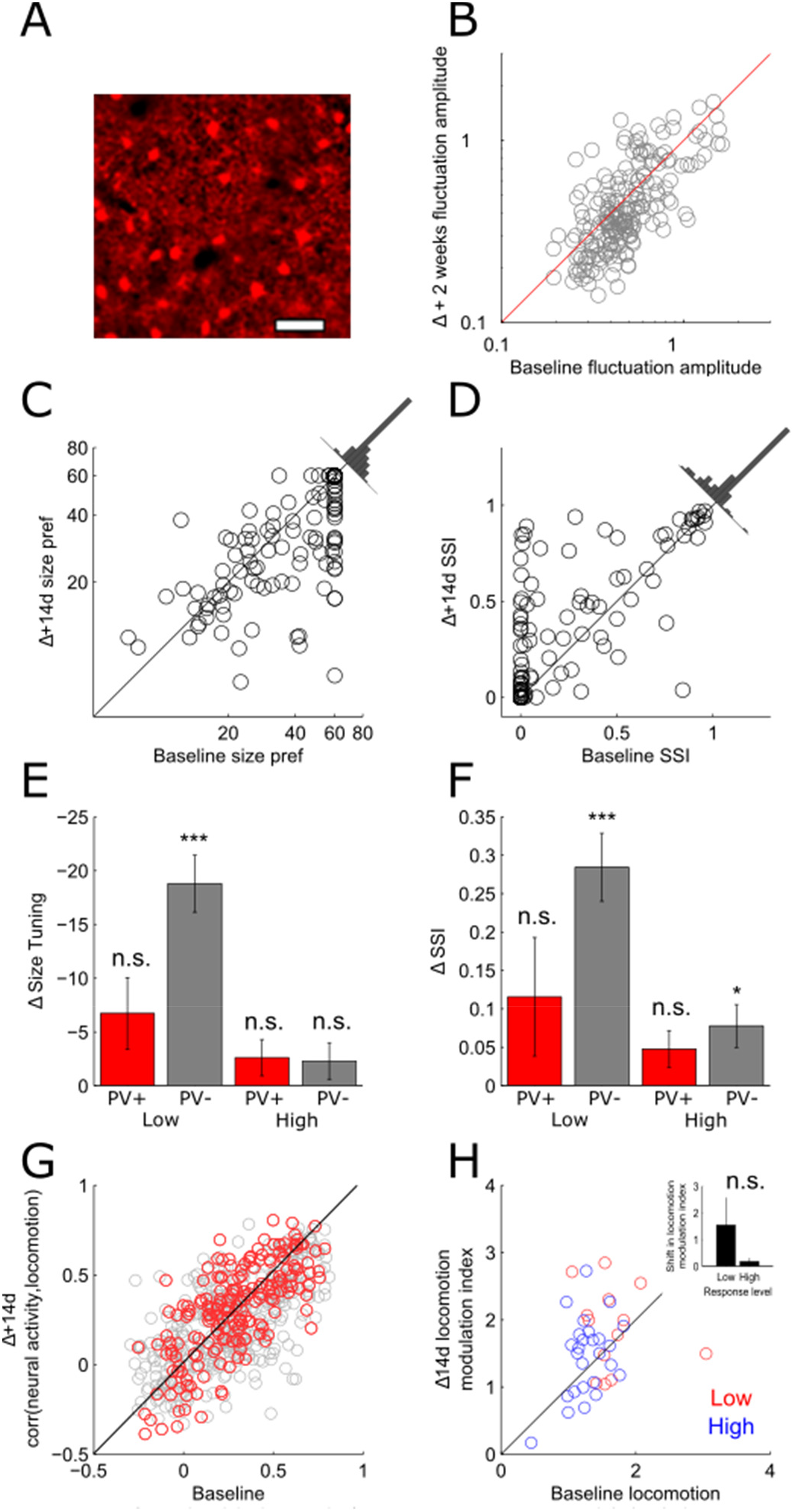
Greater stability of size preference and SSI in PV+ interneurons. (A) Example of average frame of red channel showing labelling of PV+ interneurons expressing tdTomato. (B) Intersession correlation in magnitude of calcium signal fluctuations. (C-D) Intersession correlation of size preference (C) and SSI (D). (E-F) Comparison of intersession shift in size preference (E) and SSI (F) showing greater intersessional plasticity on average in putative excitatory neurons. (G) Correlation of neural activity with locomotion between sessions in PV+ interneurons (red) and putative excitatory neurons (grey). (H) Intersession comparison of LMI in low (red) and high (blue) responding neurons. Inset bar plot shows mean shift in LMI in low and high response groups. All data are presented as mean ± SEM ***p < 0.001; **p < 0.01; *p < 0.05; n.s., not significant.

## Discussion

We used chronic 2-photon imaging of the primary V1 to make longitudinal measurements of the influence of two types of contextual factors on activity of visual cortical neurons; surround suppression and modulation of neural activity by locomotion. These results provide, to the best of our knowledge, the first longitudinal measurements of the stability of the influence of these contextual factors on visual cortical processing in awake animals. We found that a subset of highly visually responsive putative excitatory neurons maintained stable size preferences over the 14d period, while in contrast less strongly visually responsive neurons were significantly more variable between sessions. Unexpectedly this greater intersessional variability was not random, but was instead characterised by systematic shift towards smaller size preferences and greater surround suppression. This plasticity may be driven by the strong visual stimulation during the baseline experiment (as opposed to normal visual experience between imaging sessions) as the visual cortex is considered to be mature, and outside of the developmental critical period at the age studied (Gordon and Stryker, 1996; Wang et al., 2010). In contrast to the putative excitatory population as a whole, the PV+ inhibitory population exhibited a greater degree of stability and did not show strong evidence of plasticity driven by the visual stimulus. Both cell classes showed a range of degrees of modulation of activity by locomotion behaviour, but a high degree of between-session stability, such that highly locomotion modulated neurons in the baseline session continued to be highly locomotion modulated 14d later. In addition we found evidence in low responding neurons in both cells classes of an increase LMI between sessions although this was only found to be robust and statistically significant in putative excitatory neurons.

An outstanding question regards the means by which highly responsive excitatory neurons attain their greater stability. In additional to exhibiting a greater degree of stability, highly responsive excitatory cells in the majority of cases preferred relatively small stimuli and were subject to a high degree of surround suppression. Given that a large fraction of the intersessional variability in more weakly responding neurons appears to be accounted for by systematic stimulus driven reductions in preferred size and SSI, one explanation could be a ceiling effect whereby highly responsive cells already have as small a size and as high degree of suppression as is achievable. While this may contribute to the greater stability of some cells, this does not appear to be a complete explanation as there are a large fraction of highly responsive cells stably tuned to intermediate sizes. A second explanation could be that the poorer signal to noise ratio of recordings from more weakly responding cells could result in greater inter-session variability simply due to greater measurement noise. Again we believe this is unlikely to be a dominant factor as between session variability was not random in weakly responding cells, but was instead almost invariably biased towards smaller size preferences and greater SSI. In addition within session analysis showed no significant shift in preferred size or SSI (Supplementary Figure 2).

Recent studies have increasingly established that a previously unexpected degree of plasticity exists in adult sensory cortex (Frenkel et al., 2006; Fu et al., 2015; Sato and Stryker, 2008). The present study further advances our understanding of adult plasticity by demonstrating that surround suppression is a highly plastic receptive field property in adult V1. In the present experiments plasticity appears to be driven by the high contrast gratings which are shown to the animal during the baseline recording session as naturally occurring developmental maturational processes are considered to be complete by this age (Gordon and Stryker, 1996; Pecka et al., 2014; Wang et al., 2010). One intriguing possibility is that suppression is homeostatically regulated such that high levels of activity, as driven by the strong experimental stimulus, may result in upregulation of suppression. This begs the question of whether the contrary may also be true, that reduced input scales down suppression and thus whether similar processes are engaged during other plasticity inducing paradigms such as monocular deprivation. Indeed reduction of visual activity by acute dark rearing has previously been shown to enhance visual cortical plasticity, and this may occur in part through a reduction of suppressive effects. Another possibility is that the experimental stimulus induces an LTP like state (Frenkel et al., 2006) at synapses providing suppressive inputs. Either way this data highlights that even under ‘passive’ viewing conditions plasticity occurs in surround suppression, and prompts the question of the behavioural and perceptual consequences of this.

Previous studies have provided abundant evidence that visual cortical activity is modulated by locomotion with a number of circuit mechanisms proposed including input from motor cortex (Keller et al., 2012), nicotinic modulated cholinergic input from the basal forebrain (Fu et al., 2014), activation of the mesencephalic locomotor region (Lee et al., 2014) and noradrenergic input from the locus coeruleus (Polack et al., 2013). Further to this we provide evidence that each neuron has to a large extent its own unique degree to which it is locomotion modulated, and this is stably maintained over at least the time period studied here. This suggests a precision and stability of this aspect of cortical circuitry which is perhaps surprising considering some of the relatively indirect mechanisms by which locomotion has been proposed to modulate V1 activity. It also prompts the question of the functional significance of some neurons maintaining greater degrees of locomotion sensitivity. In addition we find that the effect of locomotion on visually evoked activity (the gain), is also plastic specifically in low responding neurons. The coincidence of plasticity of surround suppression and locomotion induced gain provides further evidence of common circuit elements underlying the two operations (Adesnik et al., 2013; Ayaz et al., 2013; Dipoppa et al., 2016).

A final question posed by these results pertains to why the PV+ interneuron population should exhibit reduced levels of inter-session plasticity relative to the putative excitatory population of V1 neurons? Current evidence suggests that PV interneurons receive input relatively indiscriminately from local pyramidal neurons which is thought to account in part for their broad orientation tuning (Cossell et al., 2015). We observed that contrary to a model of broad sampling of the local population's size preferences, PV+ neurons exhibited a range of size preferences and SSI values. The PV+ population also did not undergo the robust shift in size preference observed in the putative excitatory population as a whole. The latter observation could be for at least two reasons. The PV+ cells may be less plastic than the putative excitatory population. Or secondly the PV+ population's functional input might simply be dominated by the more active and less plastic fraction of the putative excitatory population.

In summary we showed that a large fraction of visual cortical cells maintain a stable degree of modulation by contextual factors during visual processing over a timescale of weeks, and that this stability was particularly apparent amongst highly responsive putative excitatory and PV+ inhibitory populations of cells. In the case of modulation of activity by locomotion this indicates that precise and stable circuitry mediate these effects, and that there are subsets of neurons which are more closely and persistently involved in integrating locomotion with visual information. Despite this stability we provide evidence that strong visual stimulation might itself be able to drive long-term increases in suppressive effects and LMI in a more labile and weakly responding subpopulation of excitatory neurons, suggesting a hitherto unexpected degree of plasticity in adult cortex of this receptive field property.

## Experimental procedures

Briefly chronic awake imaging experiments were carried out in adult C57BL/6J background animals with labelling of PV interneurons achieved using a cre driver line. Neurons in V1 were labelled with GCaMP6S using an AAV with expression under the human synapsin promotor. During imaging animals were placed on a rotary encoder coupled cylindrical treadmill, and neurons were visualised through a chronically implanted cranial window using a resonant scanning 2-photon microscope. Vascular landmarks used to relocate regions of interest between sessions. Visual stimulation were presented on a calibrated LCD screen using the psychophysics toolbox. Detailed experimental procedures are available in the supplementary materials.

---

**Supplementary Figure 1.**
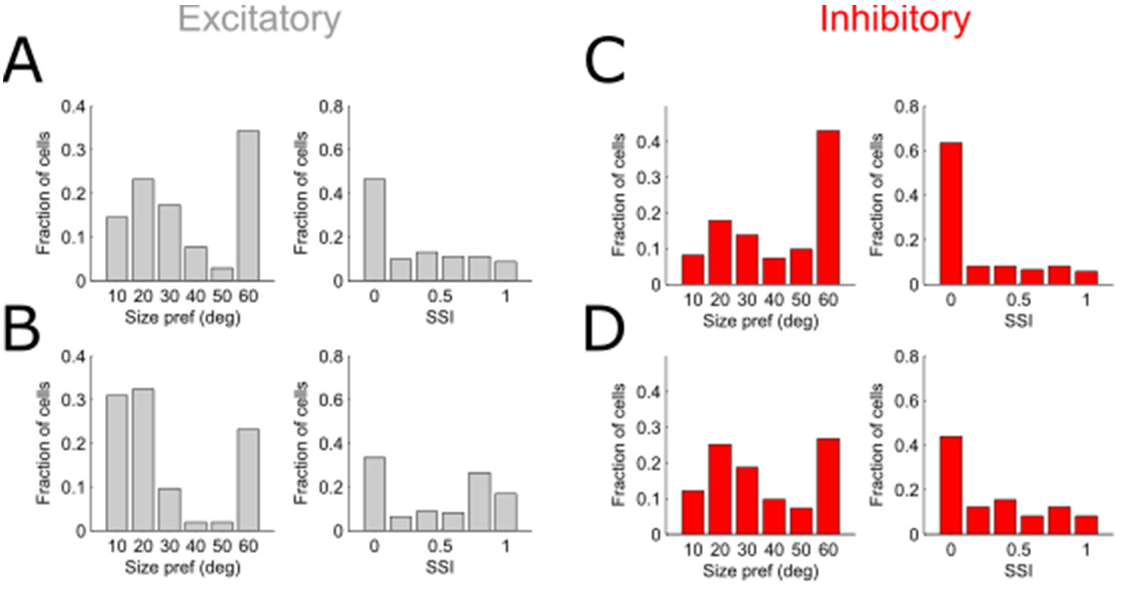
Distribution of size preferences and SSI in putative excitatory and PV+ populations. (A-D) Distribution of size preferences and SSI in putative excitatory neurons at baseline (A), putative excitatory neurons 14d later (B), PV+ neurons at baseline (C) and PV+ neurons 14d later (D).

**Supplementary Figure 2.**
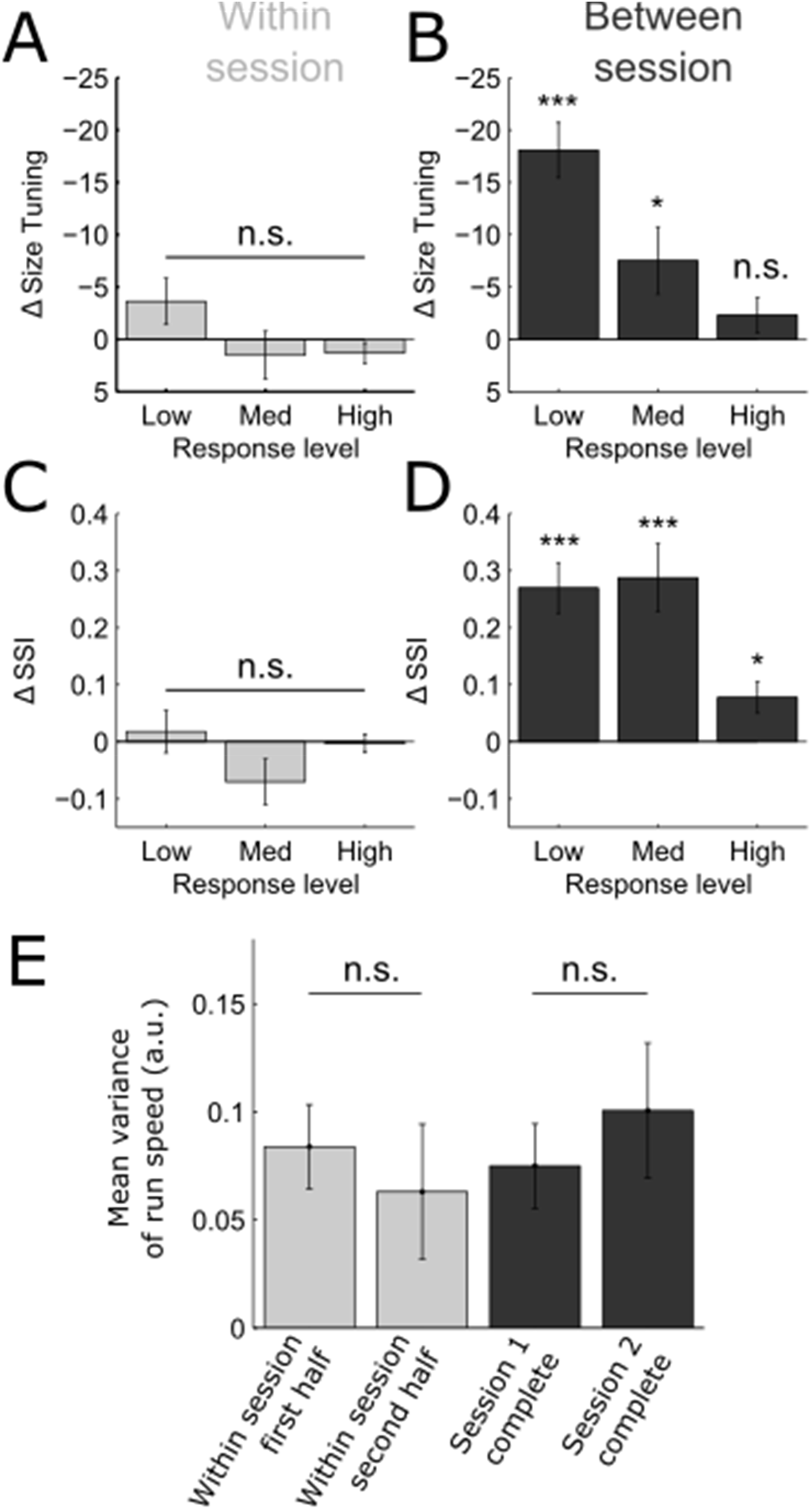
Within session verses between session stability. (A-B) Within session (A) and between session (B) stability of size tuning in the 3 response level groups. (C-D) Within session (C) and between session (D) comparison of SSI in the 3 response level groups. (E) Mean variance in running speed within the first and second half of each session (light grey) and in the first and second session as a whole (dark grey). All data are presented as mean ± SEM ***p < 0.001; **p < 0.01; *p < 0.05; n.s., not significant.

**Supplementary Figure 3.**
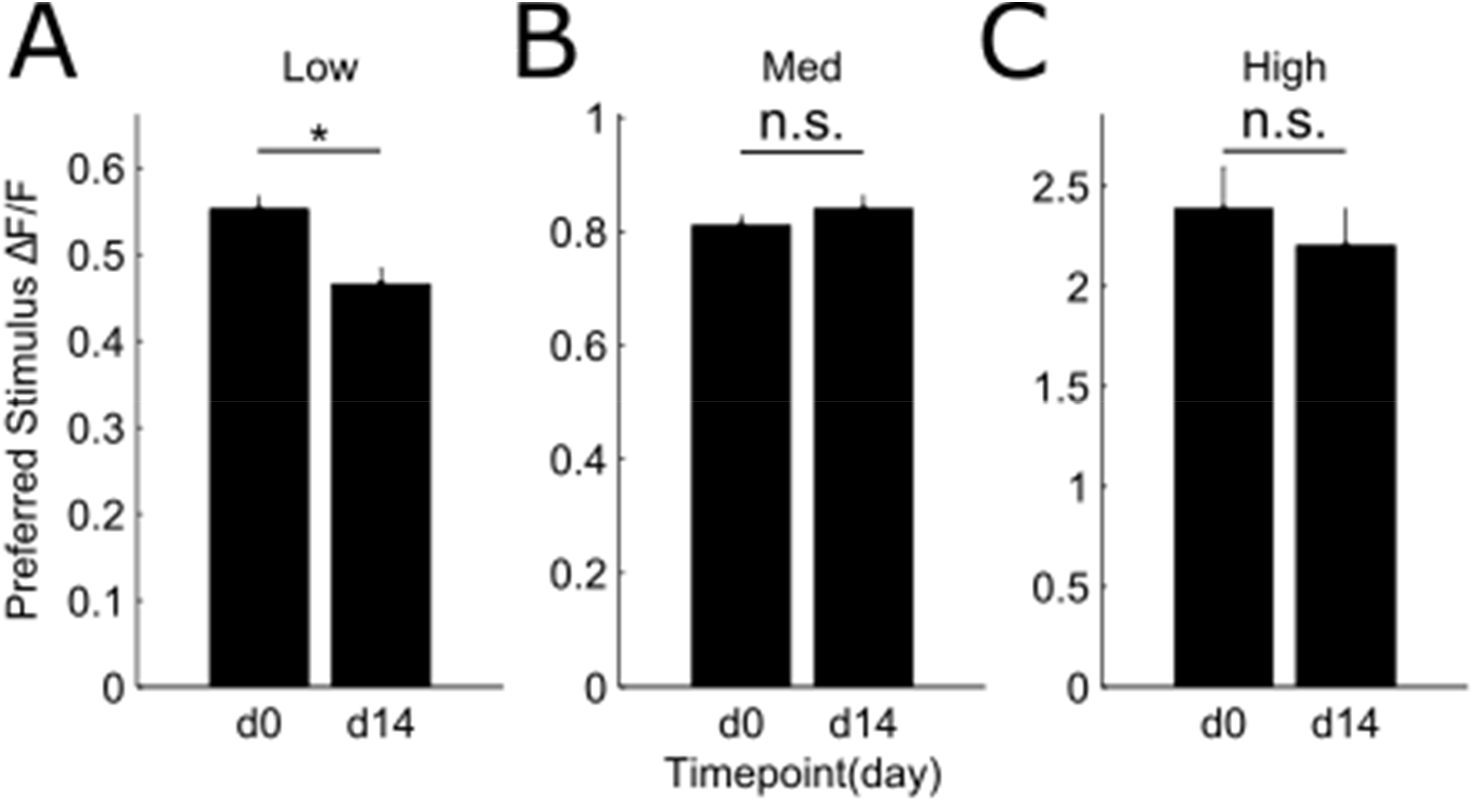
Between session response amplitudes to preferred stimuli. (A-C) Comparison of response amplitude to preferred stimulus at day 0 and day 14 in the 3 response level groups. All data are presented as mean ± SEM *p < 0.05; n.s., not significant.

**Supplementary Figure 4.**
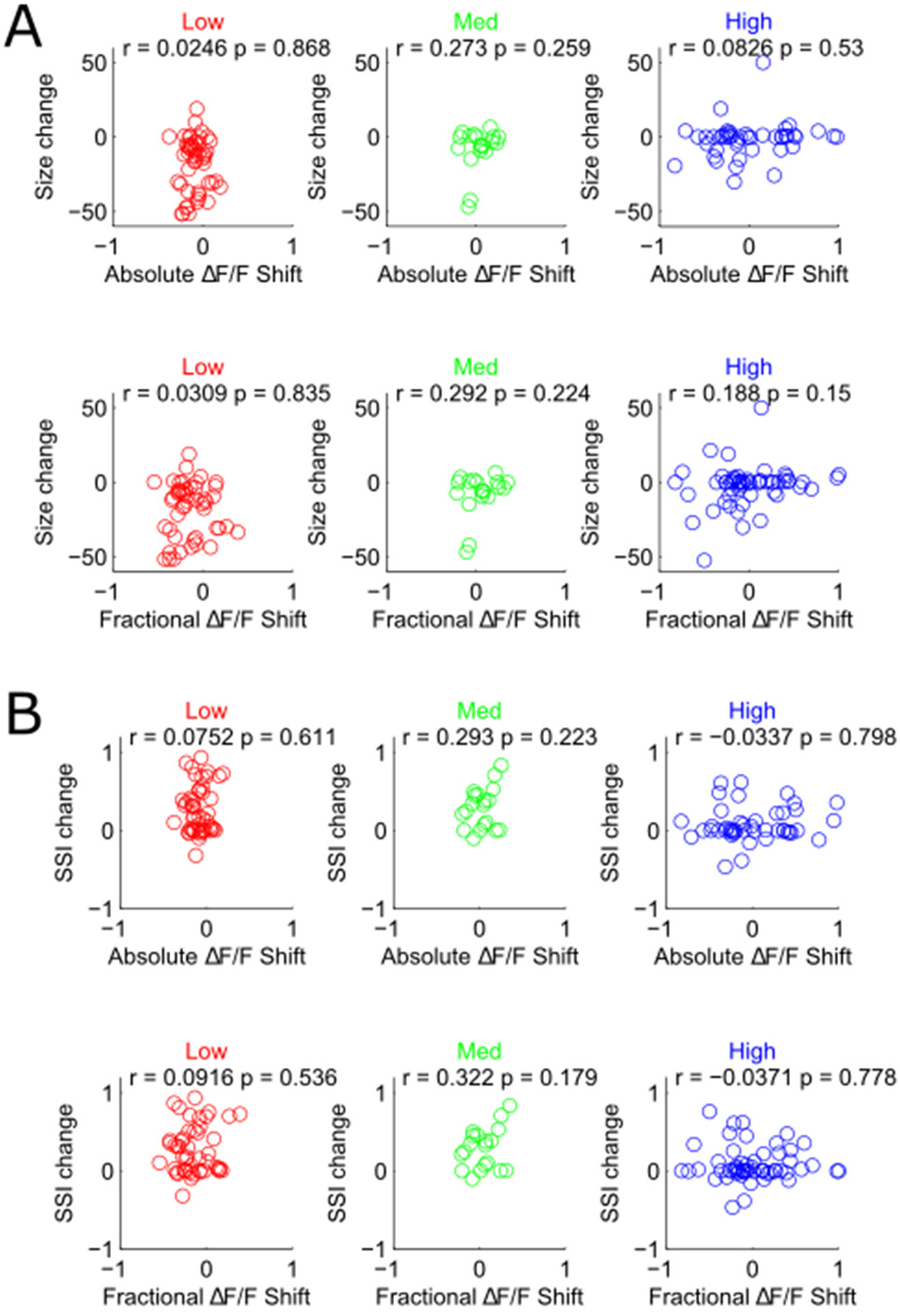
Relationship between shift in response amplitude and shift in size preference and SSI. (A) Correlation between shift in size preference and absolute shift in response amplitude (top row), and relative shift in response amplitude (bottom row), in the 3 response amplitude groups. (B) Correlation between shift in SSI and absolute shift in response amplitude (top row), and relative shift in response amplitude (bottom row), in the 3 response amplitude groups.

**Supplementary Figure 5.**
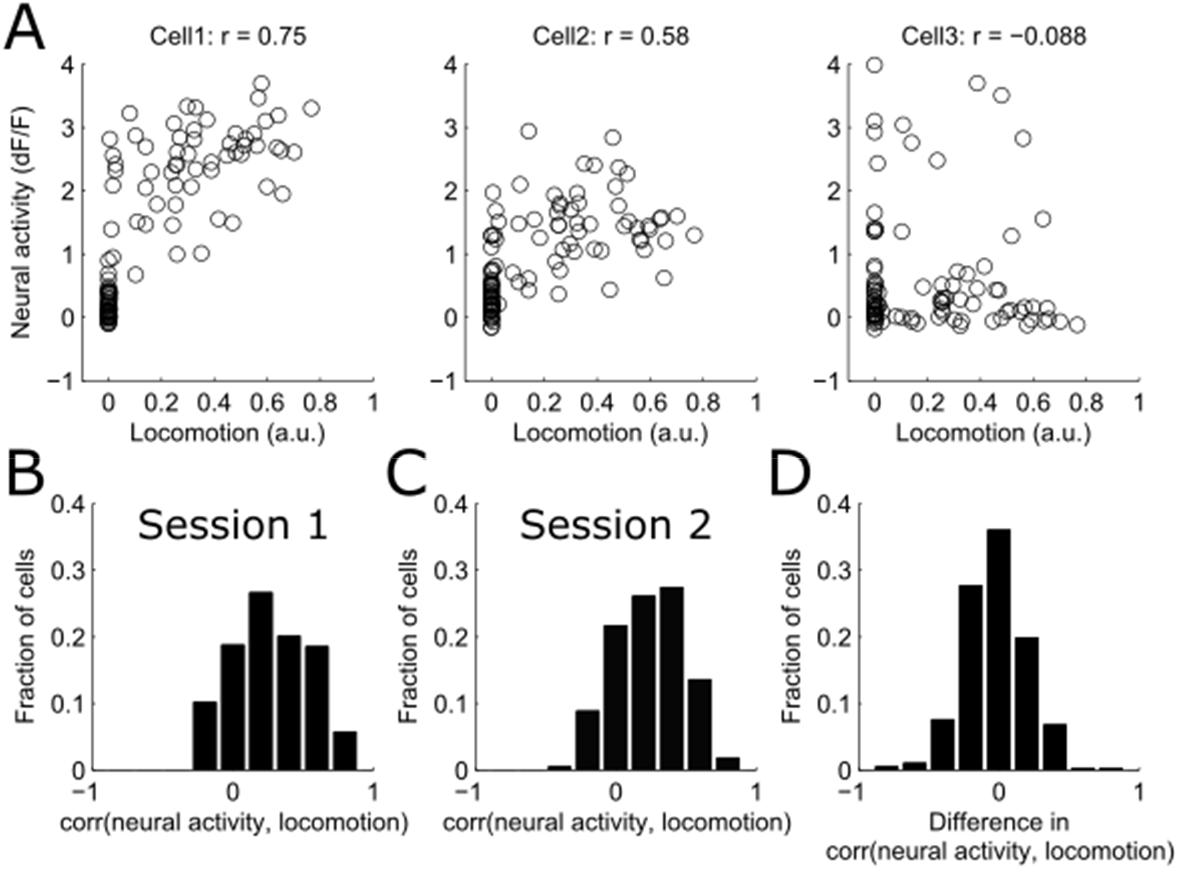
Correlations of neural activity with locomotion. (A) Examples of correlation between neural activity and locomotion in 3 neurons. Data points represent moments in time at which neural activity and locomotion velocity were sampled for individual cells. (B) Distribution of correlation coefficients in session 1 (B) and 2 (C), and distribution of differences in correlation coefficients between sessions (D).

**Supplementary Figure 6.**
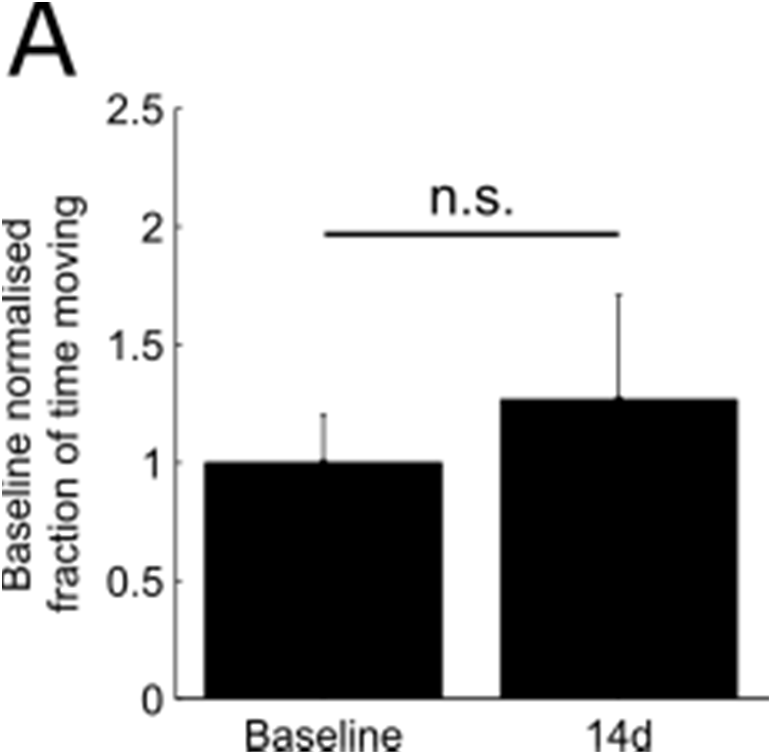
Normalized fraction of time spent moving in two sessions. n = 8. All data are presented as mean ± SEM *p < 0.05; n.s., not significant.

## Supplemental procedures

### Animals

All experimental procedures were carried out in accordance with institutional animal welfare guidelines, and licensed by the UK Home Office. Experiments were carried out on 8 adult male mice (aged P80-P95) in which PV interneurons were labelled by crossing the B6.Cg-Gt(ROSA)26Sortm14(CAG-tdTomato)Hom/J and B6;129P2-Pvalbtm1(cre)Arbr/J (Jackson Laboratory, JAX Stock#007914 and 008069 respectively). Mice were housed under normal light conditions (14h light, 10h dark) and recordings were made during the light period.

### Animal surgical preparation and virus injection

Aseptical surgical procedures were conducted based in large part on previously described protocols (Goldey et al., 2014). Approximately one hour prior to cranial window surgery and virus injection, animals were administered with the anti-biotic Baytril (5mg/kg, s.c.) and the anti-inflammatory drugs Carprofen (5mg/kg, s.c.) and Dexamethasone (0.15mg/Kg, i.m.). Anaesthesia was induced and maintained using Isoflurane at concentrations of 4%, and 1.5-2% respectively. After animals were stereotaxically secured, the scalp and periosteum were removed from the dorsal surface of the skull, and a custom head plate was attached to the cranium using dental cement (Super Bond C&B), with an aperture approximately centred over the right primary visual cortex. Transcranial intrinsic signal imaging was then used to determine the precise location of V1, after which a 3mm circular craniotomy was performed, centred on the area of V1 which responded to visual stimulation at an elevation of 20 deg and azimuth of 30-40 deg. Next, 2-3 40nl injections of a virus to drive expression of GCaMP6S (in 5 animals AAV1.Syn.GCaMP6s.WPRE.SV40; titre after dilution 2×10^11^ GC/ml; in 3 animals in which only PV neurons were labelled AAV1.Syn.flex.GCaMP6s.WPRE.SV40; titre after dilution 2×10^11^) were made into this region at a depth of 200-300µm at sites spaced by approximately 500µm. Injections were made using a microsyringe driver (WPI, UltraMicroPump) coupled to a pulled and bevelled oil filled glass micropipette with a tip outer diameter of approximately 30µm. After injection the craniotomy was closed with a glass insert constructed from 3 layers of circular no 1 thickness glass (1×5mm, 2×3mm diameter) bonded together with optical adhestive (Norland Products; catalogue no. 7106). After surgery animals were allowed at least 2 weeks in which to recover and for GCaMP6S expression to stabilise.

### Imaging and locomotor behaviour

In vivo 2-photon imaging was performed using a resonant scanning microscope (Thorlabs, B-Scope) with a 16× 0.8NA objective (Nikon). GCaMP6 and tdTomato were excited at 980nm using a Ti:sapphire laser (Coherent, Chameleon) with a maximum laser power at sample of 50mW. Data was acquired at approximately 60Hz and averaged, resulting in a framerate of approximately 10Hz. Cortical surface vascular landmarks were used to locate the same neurons between sessions. During 2-photon imaging animals were free to run on a custom designed fixed axis cylindrical treadmill, and movement was measured using a rotary encoder (Kübler, 05.2400.1122.0100). Imaging, behavioral and visual stimulation timing data were acquired using custom written DAQ code (Matlab) and a DAQ card (NI PCIe-6323, National Instruments).

In vivo intrinsic signal imaging was performed using previously described methods (Ranson et al., 2013, 2012) using either a custom built system with a MAKO G-125B camera (AVT) or a commercially available system (Imager 3001, Optical Imaging Inc.).

### Visual stimuli and experimental design

Mice were first habituated to head fixation in the experimental setup for approximately 15-30 minutes. On the following day, during the first imaging session, the preferred retinotopic location of the field of view of neurons was established using circular 30×30 deg drifting horizontal gratings in a 3×3 grid with temporal frequency of 2 Hz and spatial frequency of 0.05 cycles per degree. Each stimulus appeared and was stationary for 5 seconds, drifted for 2 seconds, was stationary for 2 further seconds and then disappeared. Trials were spaced by 3 seconds, during which a grey screen was displayed. Visual stimuli were generated using the psychophysics toolbox (Brainard, 1997), and displayed on calibrated LCD screens (Iiyama, BT481). Having established retinotopic preference, orientation tuning was next measured using circular gratings with the same temporal and spatial frequency, at the identified preferred location, and displayed at 12 different orientations. Finally size tuning was measured, again using stimuli with the same temporal and spatial frequency, and at the preferred location, with a horizontally orientated grating, with stimuli sizes ranging from 10 – 60 deg in steps of 10 deg. During the second imaging session, 14d later, the same neurons were relocated using vascular landmarks and the size tuning protocol was repeated. It is important to note that while the orientatation, spatial frequency and stimulus centring will inevitably have been suboptimal for many neurons, this was constant between sessions and thus any underestimate of size preference or SSI would have effected both sessions similarly.

### Calcium imaging and behavioural data analysis

Brain motion was first corrected for using an automated registration algorithm (Guizar-Sicairos et al., 2008) implemented in Matlab, and data from the second imaging session was registered to the first. A 20µm border was removed from all frames (more than the maximum brain movement observed) to ensure that all pixels were present in all frames in both sessions. Regions of interest were next identified from data acquired during the first session using custom written a semi-automated algorithm based on grouping of pixels with correlated time-courses. Pixels within each region of interest were averaged and background fluorescence contamination was estimated from a 30µm circular area surrounding each soma ROI (excluding other somas) and subtracted from the soma ROI signal with a weighting of 0.7. Only cells with somas which were >5% brighter than surrounding neuropil were included in further analysis. The time series of each ROI was then converted from a raw fluorescence value to ΔF/F with the denominator F value calculated by smoothing the trace and then calculating a sliding window minimum with a window size of approximately 20 seconds. Cells were semi-automatically classified as PV+ based on a thresholded mean registered red channel image, which was eroded and then dilated to remove small areas of labelling of neural processes.

The average responses of neurons to each visual stimulus was calculated, and the max ΔF/F value during the 2s drifting phase was taken as the neuron's response amplitude. Unless the effect of locomotion was being explicitly analysed visual stimulus trials in which animals were moving were excluded. Orientation tuning data were fit using a sum of two Gaussians with identical widths which were constrained such that one peaked at the preferred stimulus, and the peaks were 180 degrees apart (Carandini and Ferster, 2000):

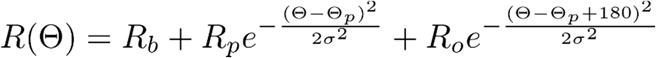

Here, R(θ) is the response to the orientation θ, R_b_ is the baseline response, R_p_ is the response to the preferred orientation, R_o_ is the response to the preferred orientation but with the opposite direction of motion, σ is the tuning width, and brackets indicate orientation values between 0° and 180°.

Size tuning data was fit using a Difference-Of-Gaussians model (DeAngelis et al., 1994):

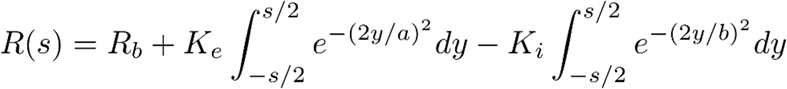

Here, *R*(*s*) is the response to size *s*, *R_b_* is the baseline response, and the two integrals represent excitatory and inhibitory components.*K_e_* and *K_i_* are the gain of the excitatory and inhibitory components, and *a* and *b* represent space constants. Surround suppression index (SSI) was defined as (R_pref_-R_max_)/R_pref_ where R_pref_ is the response to the preferred fitted size and R_max_ is the fitted response to the largest size of stimulus. Parameters were fit using the EzyFit Matlab toolbox by a nonlinear minimisation of the sum of squared residuals.

In all experiments animals spent more time stationary than moving. In order to correlate locomotion to neural activity, without introducing artefacts due to differences in fraction of time moving, all locomotion/neural activity samples were selected during which the animal was moving (i.e. m samples, which make up the ‘moving set’), and then 100 sets of m locomotion/neural activity samples were randomly selected during which the animal was stationary (resulting in 100 ‘stationary sets’ of m samples). Each of the ‘stationary’ locomotion/neural activity sets was concatenated with the ‘moving’ locomotion/neural activity set, resulting in 100 sets of moving locomotion/neural activity data (each with 2*m samples) composed of 50% stationary and 50% moving periods. The corr(neural activity, locomotion) was then calculated for each of the 100 composite sets, and the average correlation coefficient was calculated. In order to compare variation in locomotion within session verses between session (Supplementary Figure 2E), a 10 sec running average was calculated of running speed for each session, mean variance was then calculated of this running average, either in the first and second half of each session, or in the complete first and second session.

Pixel-wise stimulus preference maps were constructed by first calculating the mean of the registered imaging frames recorded during the drifting phase of each stimulus, and then determining for each pixel the stimulus which elicited the largest mean response.

## Author contributions

A.R. designed, conducted and analysed experiments, and wrote the manuscript.

## Acknowledgements

For the use of GCaMP6S we acknowledge Vivek Jayaraman, Rex A. Kerr, Douglas S. Kim, Loren L. Looger, Karel Svoboda from the GENIE Project, Janelia Farm Research Campus, Howard Hughes Medical Institute. We thank Chris Burgess for developing parts of the data acquisition system, and Kenneth Harris for developing the ROI detection algorithm. We also thank Frank Sengpiel and Kenneth Harris for helpful discussion and comments on the manuscript. This work was supported by a Wellcome Trust ISSF Seedcorn Grant to A.R., and a Wellcome Trust Strategic Award (503147).

## Highlights

- Highly active excitatory neurons are stably modulated by visual context
- Lower-activity neurons exhibit plasticity of influence of visual context in mature V1
- PV+ interneurons maintain relatively stable modulation by visual and behavioural context
- Majority of excitatory neurons are stably modulated by behavioural context

## ETOC

Using chronic in vivo 2-photon calcium imaging, Ranson et al show that while subsets of highly active excitatory neurons and PV+ interneurons are stably influenced by contextual factors, less active neurons are subject to experience dependent plasticity of these influences.

